# SCelVis: Powerful explorative single cell data analysis on the desktop and in the cloud

**DOI:** 10.1101/713008

**Authors:** Benedikt Obermayer, Manuel Holtgrewe, Mikko Nieminen, Clemens Messerschmidt, Dieter Beule

## Abstract

**Background:** Single cell omics technologies present unique opportunities for biomedical and life sciences from lab to clinic, but the high dimensional nature of such data poses challenges for computational analysis and interpretation. Furthermore, FAIR data management as well as data privacy and security become crucial when working with clinical data, especially in cross-institutional and translational settings. Existing solutions are either bound to the desktop of one researcher or come with dependencies on vendor-specific technology for cloud storage or user authentication.

**Results:** To facilitate analysis and interpretation of single-cell data by users without bioinformatics expertise, we present SCelVis, a flexible, interactive and user-friendly app for web-based visualization of pre-processed single-cell data. Users can survey multiple interactive visualizations of their single cell expression data and cell annotation, and download raw or processed data for further offline analysis. SCelVis can be run both on the desktop and cloud systems, accepts input from local and various remote sources using standard and open protocols, and allows for hosting data in the cloud and locally.

**Methods:** SCelVis is implemented in Python using Dash by Plotly. It is available as a standalone application as a Python package, via Conda/Bioconda and as a Docker image. All components are available as open source under the permissive MIT license and are based on open standards and interfaces, enabling further development and integration with third party pipelines and analysis components. The GitHub repository is https://github.com/bihealth/scelvis.

## Introduction

Single-cell omics technologies, in particular single-cell RNA sequencing (scRNA-seq), allow for the high-throughput profiling of gene expression in thousands to millions of cells with unprecedented resolution. Recent large-scale efforts to catalogue and describe all human cell types (Regev et al., 2017) dovetail with ongoing investigations to study cells and tissues in health and disease, e.g., as proposed by the LifeTime consortium (https://lifetime-fetflagship.eu). Single-cell sequencing could therefore become a routine tool in the clinic for comprehensive assessments of molecular and physiological alterations in diseased organs as well as systemic responses, e.g., of the immune system. The enormous scale and high-dimensional nature of the resulting data presents an ongoing challenge for computational analysis (Stegle, Teichmann, & Marioni, 2015). Ever more sophisticated methods combining more conventional genomics approaches with deep learning frameworks (Eraslan, Avsec, Gagneur, & Theis, 2019) allow to overcome technical limitations and biases and extract multiple layers of information, e.g. from cell types to lineages and differentiation programs. Many of these methods, their mathematical background, and the underlying assumptions will remain opaque to users without specific bioinformatics expertise. At the same time, an in-depth understanding of cell types, their functional specialization and modification by diseases, and underlying molecular correlates is often beyond the biological know-how of typical bioinformatics researchers. More than ever, single-cell omics requires close communication and close collaboration from wet and dry lab experts. Due to the large amount of data, communication need to be based on interactive channels (e.g., web-based apps) rather than static tables. Further, as single-cell omics moves towards the clinic, FAIR (Wilkinson et al., 2016) data management, data privacy, and data security issues need to be handled appropriately. All employed methods should be able to scale towards handling a large number of users and even larger numbers of samples.

### State of the Art

Web apps have been used extensively in the single-cell literature and are most commonly built on Shiny (RStudio Inc., 2014). However, standalone and general-purpose tools are to our knowledge quite rare. Pagoda (Fan et al., 2016) comes with a simple intuitive web app, which is limited to Pagoda output and requires manual preprocessing. Cerebro (Hillje, Pelicci, & Luzi, 2019) is a Shiny web app combined with a Docker container and an Electron (https://github.com/electron/electron) standalone app and provides relatively rich functionality such as gene set enrichments and quality control statistics, but relies on extensive manual preprocessing and is not (yet) ready for larger frameworks. On the other hand, the Broad Single Cell Portal (https://portals.broadinstitute.org/single_cell) provides a large-scale web service for a large number of users and studies. It includes a 10X Genomics data processing pipeline and user authentication/account management. However, the underlying Docker image strongly depend on vendor-specific cloud systems such as Google cloud and Broad Firecloud services. Its implementation thus poses practical hurdles, in particular if it is to be integrated into existing clinical infrastructure.

## Materials & Methods

SCelVis is based on Dash by Plotly (Plotly Technologies Inc., 2015) and accepts data in HDF5 format as AnnData objects, which can be created using Scanpy (Wolf, Angerer, & Theis, 2018). It also provides conversion functionality from raw text or 10X Genomics CellRanger output. The built-in converter is accessible from the command line and a web-based user interface (Figure 1). It allows for converting pipeline output with an optional description file into a single AnnData HDF5 file. One HDF5 file or a folder containing multiple such files can then be provided to SCelVis for visualization, and data sets can be selected for exploration on the graphic web interface. To enable both local and cloud access, data can be read from the file system or remote data sources via the standard internet protocols FTP, SFTP, and HTTP(S). SCelVis also provides data access through the open source iRODS protocol (Rajasekar et al., 2010) or the widely-used Amazon S3 object storage protocol. The data sources can be given on the command line and as environment variables as is best practice for cloud deployments (Adam Wiggins, 2011). The latter allows for easy “serverless” and cloud deployments.

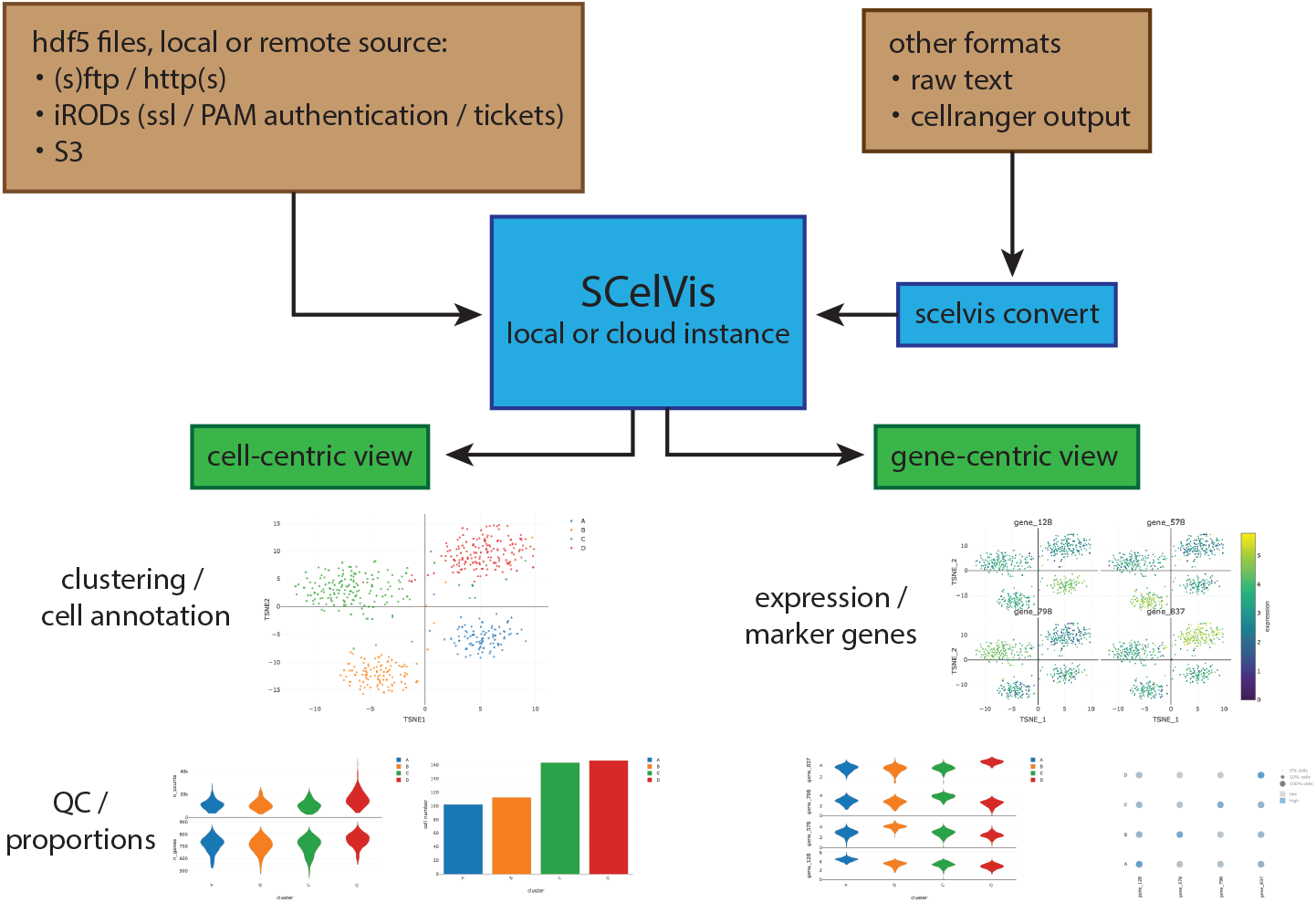

SCelVis is built around two perspectives on single-cell data (Figure 1). On the one hand, it provides a cell-based view, where users can browse and investigate cell annotations (such as cell type) and cell-specific statistics (such as sequencing depth or cell type proportions) in multiple visualizations, e.g., on a t-SNE or UMAP embedding, as violin plots or bar charts. On the other hand, it provides a gene-based view that lets users explore gene expression in multiple visualizations on embeddings or as violin or dot plots. Relevant genes can be specified by hand or selected directly from lists of marker genes.

The source code is available under the permissive MIT license on the GitHub repository at https://github.com/bihealth/scelvis. The software can be run both in the cloud and on workstation desktops via Docker.

### Usage Example

We provide two example datasets within our Github repository (see above, it also contains a link to a public demonstration instance). First, a small synthetic simulated dataset created for illustration purposes, and secondly a publicly available processed scRNA-seq dataset from 10X Genomics containing ~1000 cells of a mix of human HEK293T and murine NIH3T3 cells (Figure 2).

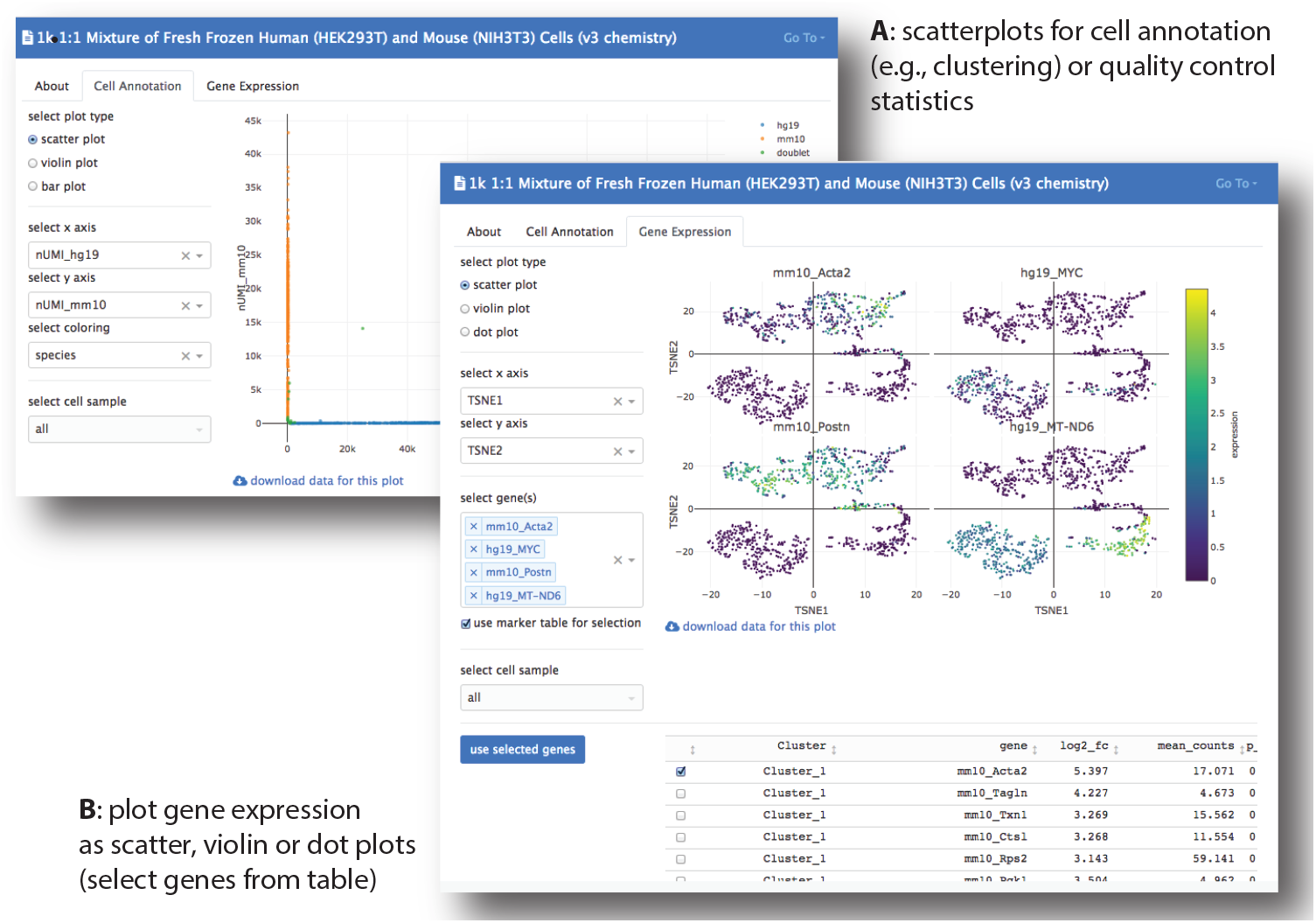

## Discussion

In this manuscript, we have presented SCelVis, a method for the interactive visualization of single-cell RNA-seq data. It provides easy-to-use yet flexible means of scRNA-seq data exploration for researchers without computational background. SCelVis takes processed data, e.g., provided by CellRanger or a bioinformatics collaboration partner, as input, and focuses solely on visualization and explorative analysis. Great care has been taken to make the method flexible in usage and deployment. It can be used both on a researcher’s desktop with minimal training yet its usage scales up to a cloud deployment. Data can be read from local file systems but also from a variety of remote data sources, e.g., via the widely deployed (S)FTP, S3, and HTTP(S) protocols. This allows for deploying it in a Docker container on “serverless” cloud systems. As both the application and data can be hosted on the network or cloud systems, the application facilitates cross-institutional research. For example, a sequencing or bioinformatics core unit can use it for giving access to non-computational collaboration partners over the internet. This is particularly interesting as it comes with no dependency on any vendor-specific technology such as the Google or Facebook authentication that appears to become pervasive in today’s life science.

## Conclusions

SCelVis is a flexible and powerful method for the visualization of single-cell RNA-seq experiments and the explorative data analysis thereof. It comes with a number of unique features, in particular complete independence of vendor-specific software or services. At the same time, it remains simple enough to be integrated as a component in more complex framework.

## Acknowledgements

The example dataset for the 1:1 mixture of human and mouse cells processed with CellRanger (v3) was taken from the 10X genomics website https://support.10xgenomics.com/single-cell-gene-expression/datasets/3.0.0/hgmm_1k_v3.

## Notes

https://github.com/bihealth/scelvis

